# Identifying residues in unfolded whole proteins with a nanopore: a theoretical model based on linear inequalities

**DOI:** 10.1101/2023.08.31.555759

**Authors:** G. Sampath

## Abstract

A theoretical model is proposed for the identification of individual amino acids (AAs) in an unfolded whole protein’s primary sequence. It is based in part on a recent report (*Nat. Biotech*. 41, 1130–1139, 2023) that describes the unfolding and translocation of whole proteins at constant speed through a biological nanopore (alpha-Hemolysin) of length 5 nm with a residue dwell time inside the pore of ∼10 μs. Here current blockade levels in the pore due to the translocating protein are assumed to be measured with a limited precision of 70 nm^3^ and a bandwidth of 20 KHz for measurement with a low-bandwidth detector. Exclusion volumes in two pores of slightly different lengths are used as a computational proxy for the blockade signal; subsequence exclusion volume differences along the protein sequence are computed from the sampled translocation signals in the two pores relatively shifted multiple times. These are then converted into a system of linear inequalities that can be solved with linear programming and related methods; residues are coarsely identified as belonging to one of 4 subsets of the 20 standard AAs. To obtain the exact identity of a residue an artifice analogous to the use of base-specific tags for DNA sequencing with a nanopore (*PNAS* 113, 5233–5238, 2016) is used. Conjugates that add volume are attached to a given AA type, this biases the set of inequalities toward the volume of the conjugated AA, from this biased set the position of occurrence of every residue of the AA type in the whole sequence is extracted. By applying this step separately to each of the 20 standard AAs the full sequence can be obtained. The procedure is illustrated with a protein in the human proteome (Uniprot id UP000005640_9606).

## 1. Introduction

Unlike DNA sequencing, for which efficient high throughput methods have become commonplace over the last two decades, a convenient easy-to-use method for protein sequencing with small sample sizes that can serve as an alternative to mass spectrometry (MS) [1,2] continues to be elusive in spite of widespread and sustained efforts. MS, while it has seen considerable success and is more or less the prevailing and preferred method, is largely a bulk method that typically requires several thousands of to possibly a million copies. The need to work with very small samples, possibly down to the single or few molecules level, arises from the large dynamic range of copy numbers of expressed proteins in the proteome, which in a single cell may be as high as 7-10 orders of magnitude, going from as few as one copy to ten million or more [3]. While MS continues to see significant decreases in sample sizes with more efficient and sensitive isolation and separation techniques [4], there has been a growing interest in alternatives to MS that can sequence single whole protein molecules.

The range of protein sequencing methods (including single molecule protein sequencing or SMPS) is quite wide, it includes nanopores, optical tags, and methods based on affinity to particular peptides or epitopes. For two recent reviews of protein sequencing see [5,6]; a feature article [7] summarizes some ongoing developments in single-cell proteomics. The following are selected examples of work within the last decade or so in protein sequencing, including NIH-funded SMPS projects in progress: 1) Nanopores designed for SMPS [8]; 2) SMPS by sequential isolation and identification of N-terminal amino acids attached to a linker molecule [9]; 3) SMPS with nanopores and Raman Spectroscopy [10]; 4) Detecting single AA substitutions in individual molecules [11]; 5) unfoldase-coupled nanopore array technology [12] and protein arrays with epitopes for large-scale peptide recognition [13]; 6) recognition tunneling [14]; 7) theoretical methods such as transverse detection in a nanopore [15] and identification of cleaved residues with a tandem electrolytic cell (e-cell) [16]; and 6) sequencing of full-length proteins using N-terminal AA binders (NAABS) [17]. Almost all of the above are based on analog measurements. A theoretical alternative with binary-digital measurements based on the superspecificity of transfer RNAs is described in [18].

Many protein sequencing methods, bulk and single molecule, use some form of proteolysis of a whole protein into peptides. The latter are sequenced and the results are combined by using pattern matching algorithms and known protein sequences to get the sequence for the whole protein. In some cases AI and machine learning methods are used [19]. Strictly these are not *de novo* methods as they rely on existing proteome databases. Alternatively the terminal residue may be identified and cleaved, or cleaved first and identified next. Successive such steps results in the sequence of the whole protein being known. Both approaches are destructive of the sample so neither reuse nor multiple measurements are possible. Whole protein sequencing in contrast is based on successively identifying residues in the protein either in the natural or some other order without any breakage. The number of known whole protein sequencing methods is small.

### The present work

A theoretical framework for whole protein sequencing based on uniform translocation at constant speed [20] through two pores of slightly different lengths (5 nm, 4.5 nm) is presented. The model includes realistic limitations imposed by detector bandwidth and available current blockade level precision. Translocation through two pores of slightly different lengths is measured with sampling pulse sequences at two slightly different bandwidths phase shifted relative to each other. From the resulting measured signals the exclusion volume of every AA in the protein sequence can be estimated to lie between two bounds within the available precision. These pairs of inequalities are used to place each residue in the whole sequence in a small subset of the AAs. To identify a residue exactly the corresponding AA volume is increased by a significant amount by attaching a high volume conjugate molecule to every occurrence of the AA in the sequence. This selectively modifies the inequalities and biases them toward the AA type so that the positions of every occurrence of the AA along the sequence can be extracted in a straightforward way. By repeating this for every one of the 20 AA types the whole protein can in principle be sequenced exactly. The procedure is illustrated with a protein from the human proteome (Uniprot id UP000005640_9606).

## 2. Polymer analysis with nanopores

Figure 1 shows a schematic of an electrolytic cell (e-cell) with a nanopore of length L in a membrane separating two chambers named *cis* and *trans* containing an electrolyte, typically KCl. When a voltage is applied across the pore, it results in an ionic current due to flow of K^+^ and Cl^-^ ions through the pore. If an analyte polymer, such as a protein molecule, is added to *cis*, it is drawn to the pore and translocates through to *trans*, leading to a decrease or blockade in the ionic current. The blockade level can in principle be used to identify the analyte or monomers in the polymer [21].

**Figure 1.**
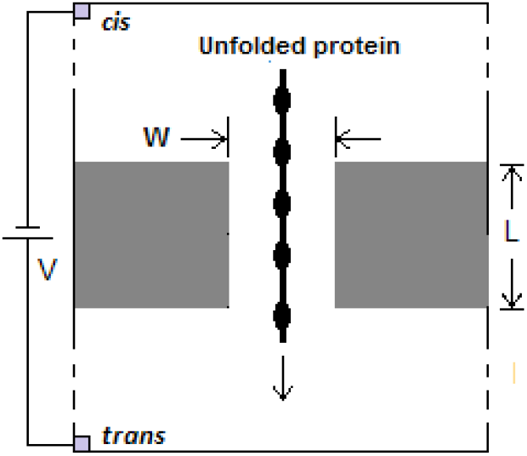
Schematic of electrolytic cell (e-cell) with nanopore in membrane separating *cis* and *trans* chambers filled with electrolyte KCl. Voltage V applied across pore ionizes electrolyte, results in electrolytic current due to Cl^-^ (K^+^) ions traveling from *cis* (*trans*) to *trans* (*cis*). Analyte (protein) molecule added to *cis* translocates through pore and causes current blockade. W = diameter of pore, small enough to cause only a single polymer strand to translocate, typically 2-4 nm; L = length of pore (= 5 or 4.5 nm in present study).

There are several advantages to nanopore sequencing of polymers compared to other methods such as MS (with proteins). Thus it requires only electrical measurements with low voltages, it may be done without labels, it can work with just one molecule or be scaled to large numbers of molecules, and it can be designed to unfold a protein prior to the latter passing through the pore. On the other hand there are also disadvantages that have hindered its early development and projected widespread use. They include: 1) the high speed with which an analyte passes through the pore with or without an applied voltage across the pore; 2) the non-uniform rate of translocation; 3) the possibility of blockage; and 4) the need to ‘thread’ the molecule into the pore from *cis* and translocate it to *trans*. A number of solutions to these problems have been proposed and/or implemented to varying extents. Much work has gone into controlling the speed of translocation of an analyte through a nanopore under the influence of diffusion and electrophoresis [22]. Slowdown methods include use of an enzyme motor [23], use of an SDS sheath [24], and a theoretical model based on a bi-level voltage profile in conjunction with pH control [25]. Most recently electro-osmosis has been used in combination with an engineered charge-selective nanopore for capture, unfolding, and translocation of polypeptides without using an enzyme motor [26].

The present work is based in part on a recent report [20] that describes a method of controlled protein translocation through the pore that unfolds the protein prior to its entry into the pore, is unidirectional and at a uniform rate, and is independent of the protein (based on experiments with a small but varied set of proteins). The method hinges on the use of guanidine chloride in the buffer, which leads to all 20 AAs having approximately the same speed of translocation through the pore and a dwell time of about 10 μs, corresponding to an exit rate of one AA into the *trans* chamber every μs. The pore used is the biological nanopore alpha-hemolysin with a length of 5 nm.

## 3. A theoretical framework for residue identification with a nanopore based on a system of linear inequalities

The model of residue identification presented is made more realistic by imposing a minimum set of constraints that can be satisfied in practice. The residue identification process is transformed into a system of linear inequalities that can be solved with well-established computational methods including but not limited to linear programming.

### 3.1 Practical constraints

In practice nanopore protein sequencing, often based on measurement of current blockade levels, regardless of the method used, needs to work with a number of limitations:

1. Limited bandwidth: One of the obstacles to reliable detection of current blockade levels due to a translocating analyte is high frequency noise, requiring large and usually impractical bandwidths. Reading protein translocation using the method in [20] would require a bandwidth of 100 KHz or more.
2. Limited precision in reading of current blockade levels: In the model presented here digitized values of analyte volume inside the pore are used as a proxy for current blockade levels. Table 1 shows the volumes (in nm^3^) of the 20 AAs. The precision used here is 70 nm^3^ and is based on [27].

**Table 1.** Volumes of the 20 standard AAs, data from [28].Volumes in nm^3^.

| AA | G | A | S | C | D | T | N | P | V | E |
| --- | --- | --- | --- | --- | --- | --- | --- | --- | --- | --- |
| Mean vol | 59.9 | 87.8 | 91.7 | 105.4 | 115.4 | 118.3 | 120.1 | 123.3 | 138.8 | 140.9 |
| AA | Q | H | M | I | L | K | R | F | Y | W |
| Mean vol | 145.1 | 156.3 | 165.2 | 166.1 | 168 | 172.7 | 188.2 | 189.7 | 191.2 | 227.9 |

### 3.2 Assumptions

1. The controlled translocation method described in [20] is assumed with an increase in the dwell time of a residue from 10 to 25 μs. A decreased bandwidth B = 20 KHz instead of the 100 KHz that would be required in [20] is assumed.
2. A profile dimension per residue (its length along the pore axis) of 0.5 nm is assumed, the same for all residues (because the residue to residue distance along the backbone of a linear stretched protein is roughly the same for all residue pairs; AA volume differences are largely in the side chain).
3. A protein is inspected by translocating through two pores of different lengths: 5 nm and 4.5 nm. Because of uniform translocation speed, at any time a pore of length 5 nm is occupied by 10 residues, while one of length 4.5 nm is occupied by 9 residues (end effects, the initial time taken to fill the pore on entry from *cis* and the final time to empty the pore as the protein exits fully into *trans*, are neglected). This difference of 1 residue in the number of residues resident in the two pores is central to the sequencing method given here.
4. The translocation signal is sampled with a pulse train with a period that is at least 1/B and a pulse width of 1/2B.
5. A volume precision of 70 nm^3^ is assumed. It is based on the study in [27] where this level of precision is used to distinguish among four subsets of the AAs that are labeled S(mall), M(edium), I(ntermediate), and L(arge) according to their volumes.

#### Notation

k = index of residue at the *trans* end of the pore

Δt = time duration of exit of *trans* end residue into *trans*

mΔt = shift of sampling pulse with respect to rising edge of first pulse in pulse train

M, M’ = length (number of residues) of protein segment inside pore (= 10 for a 5 nm pore and 9 for a 4.5 nm pore)

B = detector bandwidth in KHz

W_B_ = sampling pulse width in μs; 2W_B_ = sampling period

p_i_ = i-th residue in protein

Vol_i_ = volume of p_i_ (see Table 1)

p_k_ p_k+1_ … p_k+M-1_ = protein segment inside pore (p_k_ = residue at trans end of pore)

rVol_k,M_ = total volume of p_k_ p_k+1_ … p_k+M-1_ (there are M residues in the pore at any time)

pVol_m,B,M_ = total volume read over a pulse shifted by mΔt from origin of sampling sequence

V_step_ = volume precision = 70 nm^3^

dV_x_ = digitized volume of V_x_ = ⌊V_x_/V_step_⌋

### 3.3 Development of procedure

Consider a protein P with N residues given by

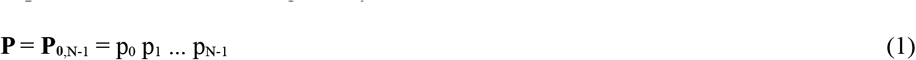

Ideally a protein can be sequenced with a nanopore that is short enough such that only one residue is inside the pore when the protein translocates through. Translocation is fast and a bandwidth in excess of 1 MHZ would be required to detect the residue as it passes through, and a very high precision would be required to distinguish between AAs whose volumes differ by very small amounts. For example, from Table 1, the difference between the volumes of M (methionine) and I (isoleucine) is only 0.9 nm^3^. With a sufficient bandwidths and volume precision the translocation signal would look something like in Figure 2, which shows the variation of residue volume along the sequence of protein number 7, id A8MUZ8, in an instance of the human proteome (Uniprot UP000005640_9606). The top panel shows the raw signal, the bottom shows residue volumes picked up by an ideal detector.

**Figure 2.**
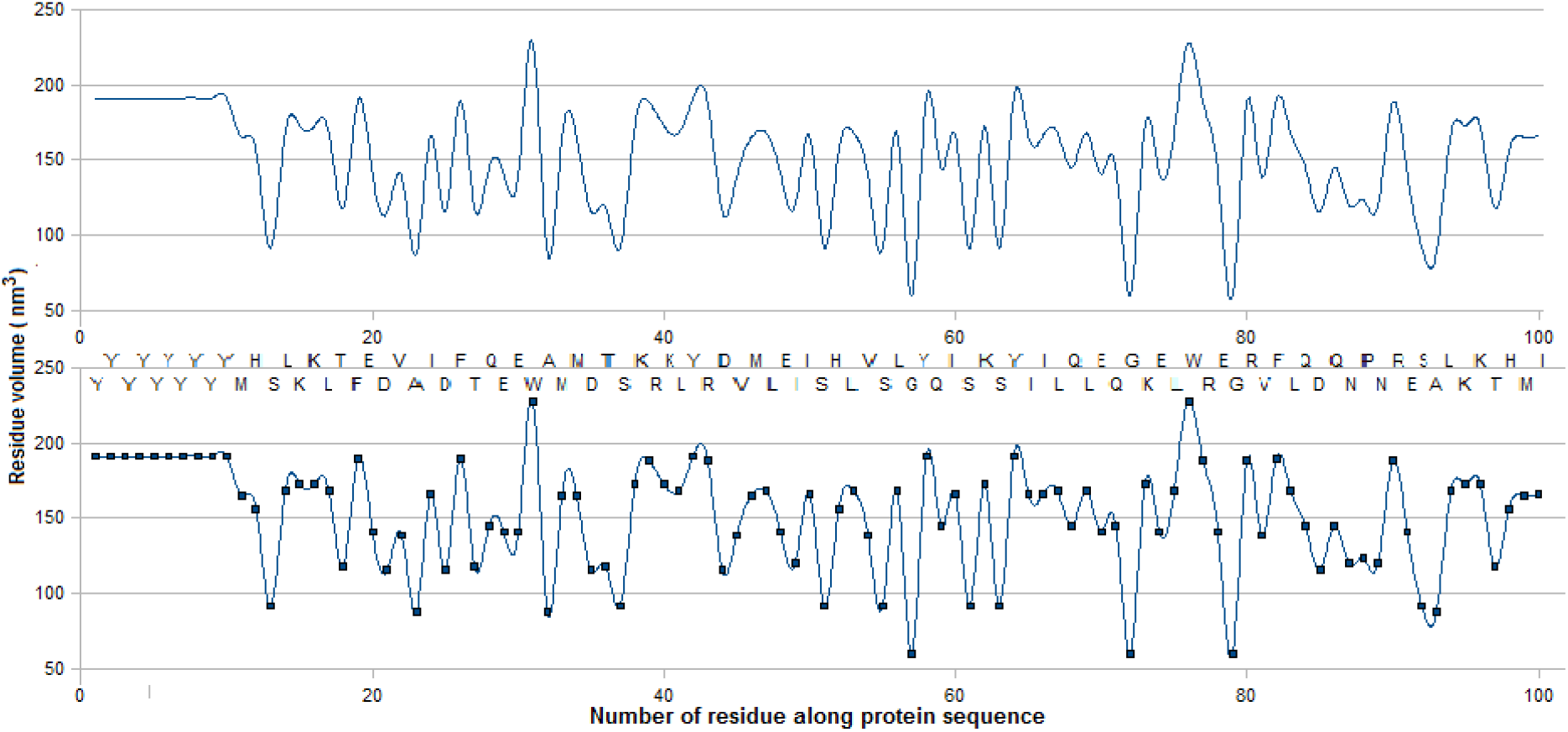
Residue volume along protein 7, id A8MUZ8, in human proteome (Uniprot UP000005640_9606) with unlimited bandwidth and volume precision

In practice none of this is possible. The assumptions in Section 3.2 are a reflection of the limitations that exist in practice. With a pore of length 5 nm, there will be 10 residues inside during the translocation process so any volume measurement is of the volume of these 10 residues. With the uniform speed of translocation assumed above, a residue exits into *trans* every μs so the resident volume keeps on changing. Figure 3 shows this changing volume, whose measurement once again is restricted by the detector bandwidth and the volume precision. Resident volumes are shown for 5 nm and 4.5 nm pores.

**Figure 3.**
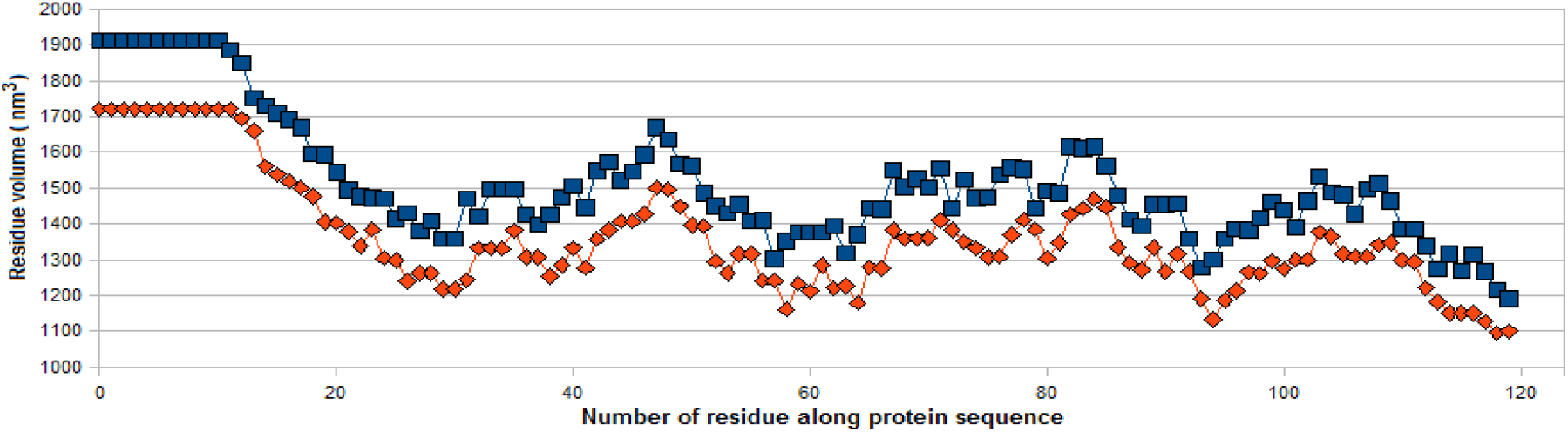
Total volume (nm^3^) of M = 10 (blue) or 9 (red) residues inside as protein translocates through pore. Number on horizontal axis corresponds to number of residue at *trans* end of pore

It is from this pore-resident volume that residues must be distinguished. A way to do this is described next. A segment with M residues from the k-th to the (k+M-1)-th that are resident in the pore is given by

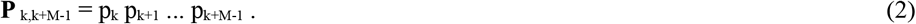

Now consider a whole protein translocating from *cis* to *trans* through the pore. Neglecting end effects for the segment in Equation 2 can be written as

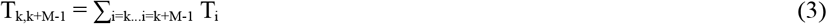

In general each AA takes a different amount of time translocating through the pore, and depending on the charge and the sign of the charge carried such translocation may not be unidirectional. As noted earlier the controlled experiments in [20] enable translocation to be uniform and unidirectional so that the dwell time for all 20 AAs is roughly the same in a 5 nm pore, about 10 μs. This corresponds to one residue exiting the pore into *trans* every Δt = 1 μs.

Now consider a whole protein translocating from *cis* to *trans* through the pore. The time taken for the segment inside the pore **P** _k,k+M-1_ = p_k_ p_k+1_ … p_k+M-1_, k=0,1,…N-M+1 to translocate out can be considered in terms of the times taken by each residue as it exits the pore. Since the latter is the same for all residues this time can be written as

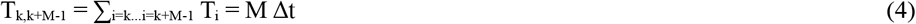

The volume excluded by this segment is

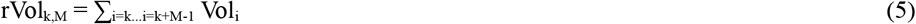

where Vol_i_ is the volume of residue i. Residue k is the next residue to exit into *trans*, so that in the next round k becomes k+1 and Vol_k_ is replaced with Vol_k+1_ and residue k+M enters the pore. This can be written as a difference equation:

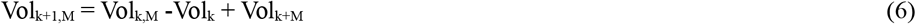

With a residue exit rate of one every μs, this value during that μs cannot be measured with a bandwidth B = 20 KHz.

Now consider the total volume of the protein sequence that moves through the pore over a sampling pulse of width 25 μs shifted through mΔt from the origin of the sampling pulse train. During this sample pulse the volume measured is given by

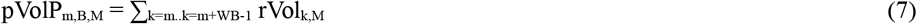

Given the limited volume measurement precision of V_step_ (= 70 nm^3^, see above), the measured volume is the digitized value

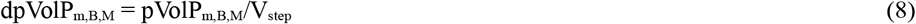

Next consider measuring a similar total volume with a sampling pulse train with period 1/2B - 1 μs = 24 μs. Over the shorter pulse width (shorter by 1 μs) shifted by 1 μs

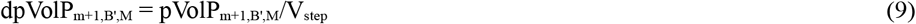

Subtracting Equation 15 from Equation 17 gives

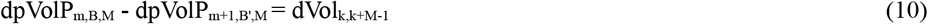

This is the digitized volume of the protein segment that is inside the pore with first residue in the segment being p_m_.

Now consider repeating the above with a pore of length 4.5 nm (which holds M’ = 9 residues, compared to M =10 with a pore of length 5 nm). This leads to

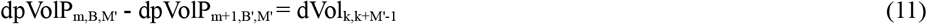

Subtracting Equation 11 from Equation 10 gives an estimate for the digitized volume of the m-th residue:

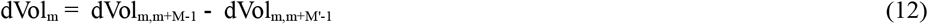

Vol_m_, the estimated volume of P_m_, satisfies

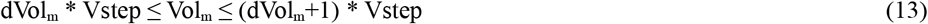

Figure 4 shows the minimum and maximum digitized volumes of residues 0 through 110 in the sample protein. Table 2 shows the estimated volumes of the 20 AAs based on these data.

**Table 2.**
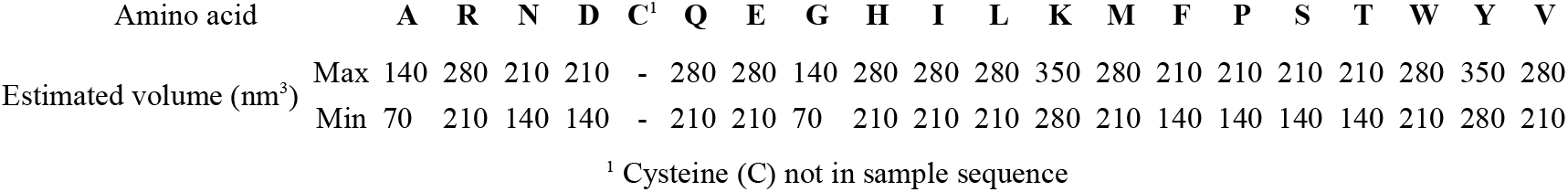
Estimated volume range for 19 of the 20 AAs.

**Figure 4.**
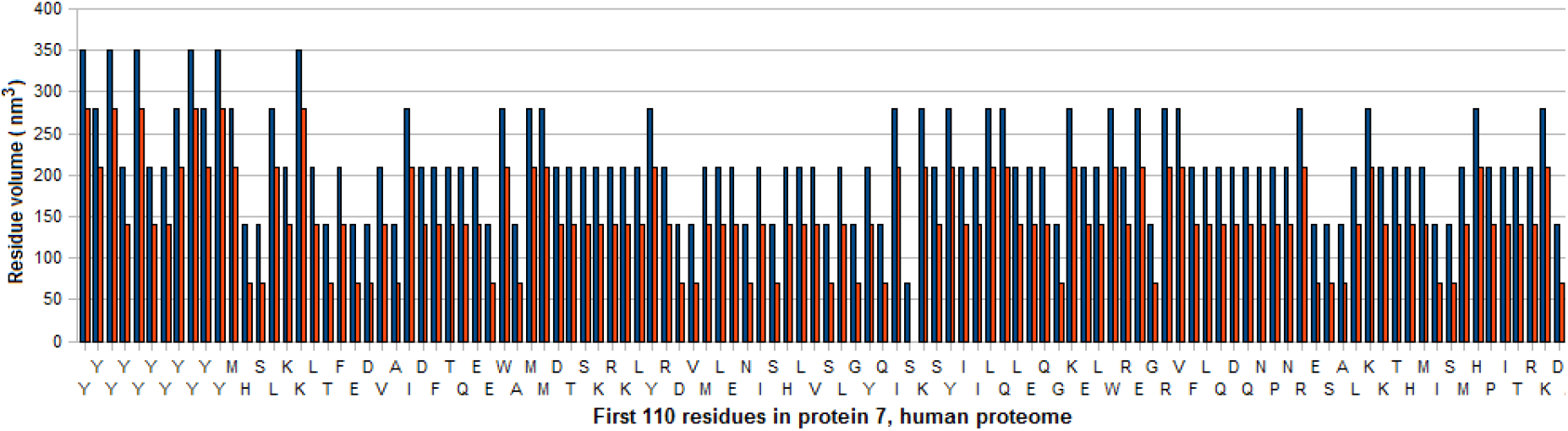
Computed bounds for volumes of first 110 residues in sample protein

Based on the bounds in Table 2, the AAs (except for Cysteine) can be divided into four subsets:

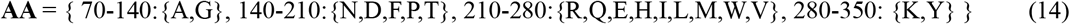

This is too coarse a division; a method for exact identification based on calculated residue volumes is given in Section 3.5 below.

### 3.4 Solution using Linear Programming methods

Solving a system of linear inequalities is equivalent to solving a Linear Programming (LP) problem [29-31]. In the present case a number of additional inequalities have to be added to get a bounded solution. The problem of determining Vol_m_ for 0 ≤ m < N can be stated as follows:

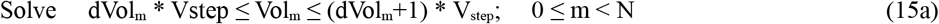

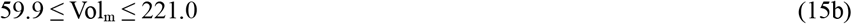

where the two bounds in Equation 15 correspond to the volumes of Glycine (G) and Tryptophan (W), the smallest and largest volume amino acids (Table 1).

Several software packages, such as lpSolve [32] are available for this. However most of the algorithms used in these packages add a dummy objective function to the given set of inequalities and are usually designed to find an extremal point in the linear polytope that is defined by a system of linear inequalities, such as Equation 15. Computational experiments on lpSolve show that these solutions are considerably off the mark and are largely unusable. In the present case it is better to use simple heuristics directly based on the bounds in Equation 13, as was done to get the results shown in Table 2.

### 3.5 Exact identification of residues

In [33] heavy tag molecules that are base-specific are attached to the bases in a DNA sequence. The DNA strand is pulled with an enzyme motor attached to the membrane near the entrance to the pore. A tag is removed by a chemical reaction when the base comes near the pore entrance. The released tag falls through the pore causing a current blockade of a unique size. As the strand ratchets through the motor the tags of successive bases are detached and identified when they fall through the pore so that the sequence of blockade levels caused by them unambiguously yields the base sequence of the strand.

A similar method can be used to identify all the instances of a given AA in the whole protein. Prior to entering the *cis* chamber for sequencing, a large volume tag molecule is attached to residues of type AA. When the translocating signal for a whole protein sequence is generated as described in Section 3.3, the sequence of digitized volumes calculated for each residue is biased toward residues with the added volume tag. As an example a tag with extra volume 500 nm^3^ is assumed to be attached to all Alanine (A) residues. Table 1 is now changed to

Figure 5 shows the digitized volumes of the residue as calculated with Equations 15 through 19 in terms of their spreads. There are four occurrences of Alanine in the sequence, their digitized volume spreads are (560,630) or (630,700), notice the clear separation of all these occurrences from those of all other AAs in the sequence.

**Figure 5.**
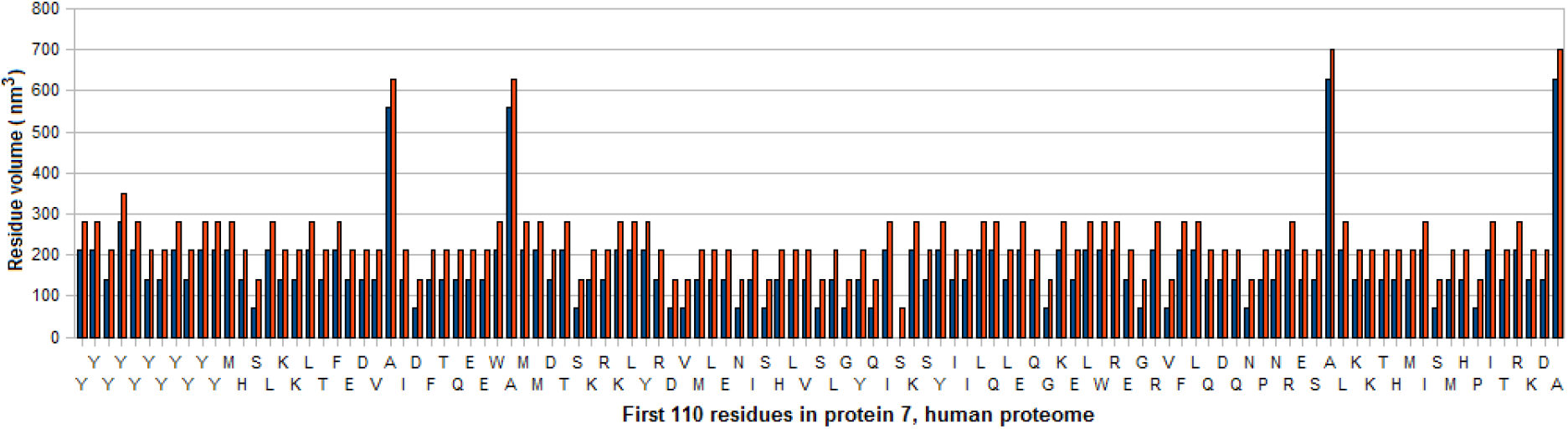
Estimated volumes of residues among the first 110 residues in sample protein with high-volume tags attached to all Alanine (A) residues

Compare with Figure 4, where the four instances of Alanine cannot be distinguished from those of Glycine (G). The ambiguity is more stark with R, Q, E, H, I, L, M, W, and V: see Equation 14.

This process can be repeated for each of the 20 AAs to get the complete sequence. In principle this is an error-free procedure.

**Table 3.**
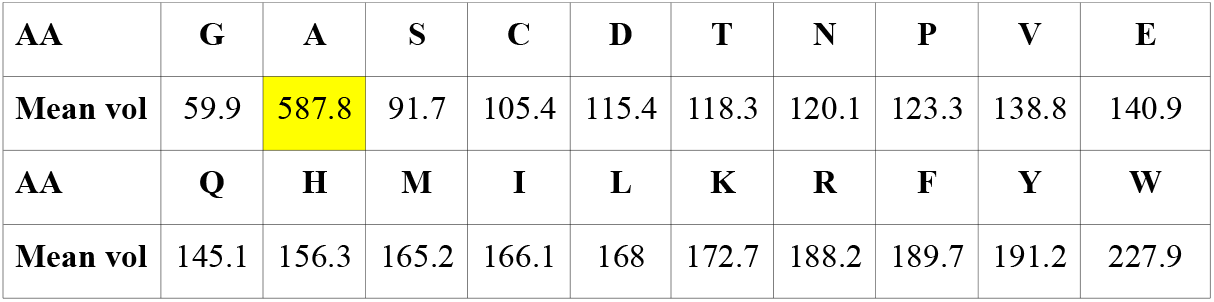
Modified version of Table 1 with volume of A augmented by attached tag with volume of 500 nm^3^. All volumes in nm^3^.

## 4 Discussion

1. The procedure given above requires shifting the sampling pulse train by 1 to W_B_ μs. This requires synchronization of the translocation signal with the sampling pulse train. Such synchronization can be done in practice by using a long homopeptide as a header with a high volume AA such as W (tryptophan) or Y (tyrosine) (see Figure 6). With a header length of 40-50 W or Y homopeptide residues (W_40_ or Y_40_), the position of the header with respect to the start of the sampling pulse can be determined from the values of measured resident volume (see Figure 3). Additionally the position of the first methionine (M) residue can be located in a straightforward way from the change in the total volume following the homopeptide header (Equations 12 and 13). If the protein were to enter the pore from the C-end the length of the trailer can be used as a resolvent; see Item 2 next.
2. The use of a header can be supplemented with a trailer consisting of a homopeptide with a different AA to allow determination of the direction of translocation- C-end to N-end or N-end to C-end. (In [20] the protein is forced to enter the pore from the N-end by attaching a homopeptide header D_n_ (n Aspartic Acid (D) residues). In the present case this D_n_ prolog can be attached to the header in Figure 6 as a charged header. (D is negatively charged and is drawn to the pore by the voltage across the pore which goes from negative on the *cis* side to positive on the *trans* side; see Figure 1.)
3. The use of high volume tags to pick out all occurrences of a tagged AA makes this a label-based method, which involves additional steps for the labeling and dissociation of a label after measurement.
4. Multiple tags with different volumes may be used to identify more than one AA in one cycle. Thus with four tags with different distinguishable volumes, only 5 cycles instead of 20 will be required. This would require the four added volumes to have sufficient separation.
5. In practice if multiple copies of the protein are available the method can be efficiently implemented in parallel. With only a single copy available the stages in Section 3.3 have to be executed in series, although many of the steps can be implemented with parallel hardware. However tagging and untagging can be a tedious step that may require a considerable amount of time.

**Figure 6.**
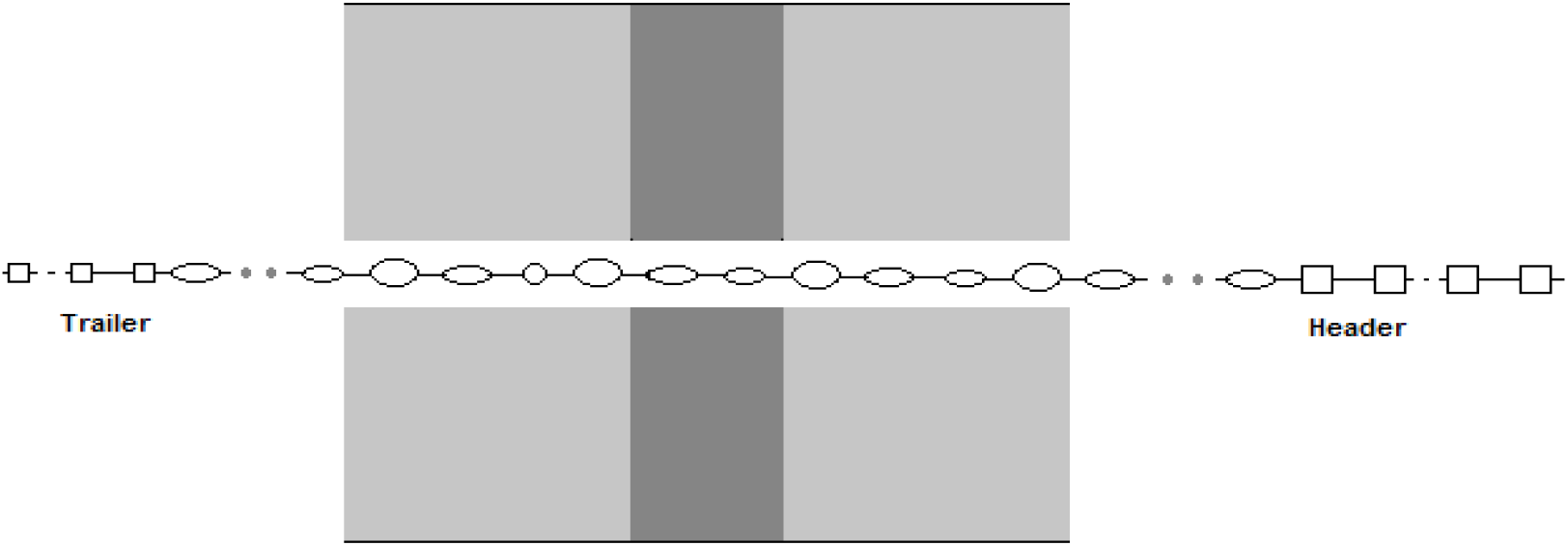
Translocation of protein with homopeptide header and trailer attached. Header and trailer may be of different AA types and different lengths for use in determining direction of entry (C-end to N-end or the reverse)

